# Camus: a video-based framework for continuous behavioral monitoring in the mouse home-cage

**DOI:** 10.64898/2026.10.23.761437

**Authors:** Tushya Mehta, Nicholas Angulo, Andrew Lu, Silas Staten-Lusty, Vincent Im, Riya Palkar, Cyrus Yorito Nadizadeh, Grace Steward, Camellia Neda Nakhjiri, Ylang Guo, Eyal Y. Kimchi

**Affiliations:** Department of Neurology, Feinberg School of Medicine, Northwestern University, Chicago IL; Northwestern University Interdepartmental Neuroscience Program (NUIN), Evanston IL

## Abstract

Home-cage activity is a window into the spontaneous behavioral repertoire of mice. These behaviors are organized across the circadian cycle, and disruptions in their duration, frequency, or organization may provide sensitive measures of neurological or psychiatric dysfunction. While continuous monitoring can help track the dynamics of spontaneous behavior, video recording in conventional laboratory cages is limited by compact cage geometry, opaque cage tops, and obstructed views of animals, food, and water. We therefore developed ***Camus***, a Camera for home-cage recordings of Mus musculus mice. Camus integrates off-the-shelf USB cameras with infrared illumination and custom 3D-printed components for access to food and water without obstructed views. The modular design pairs continuous video acquisition with a manually validated computer-vision model to quantify locomotion, feeding, drinking, and nesting-related behavior across multiday recordings. A single desktop computer can record as many as 20 cages in parallel at an incremental cost of approximately $250 per cage, recording more than 99% of expected frames at 640x360-pixel resolution and 30 frames per second. Camus captured circadian structure in locomotion, feeding, and drinking. Camus also captured nest-building and sex-specific differences in nest dynamics. Using Camus with transgenic 5xFAD mice overexpressing amyloid revealed selective, age-progressive deficits in nesting behavior. Camus provides a scalable, open-source platform that combines accessible hardware, automated video monitoring, and machine-learning based analysis for continuous and potentially longitudinal behavioral phenotyping.

## Introduction

Behavioral analysis of model systems is one of the most widely used techniques in neuroscience to understand development and aging, develop novel therapies, and discover the neural processes that shape organisms.^1–5^ Traditionally, behavioral experiments in mice involved researchers observing behavior visually.^6^ Ideal observation of mouse behavior would be continuous, longitudinal, and bias-free, using experienced observers. However, practical constraints often limit observation to only brief bouts of assessment by less experienced observers. When continuous, long-term monitoring is desired, specialized environments are often required.^7–12^ Previous attempts to collect continuous behavioral data have included the use of infrared (IR) movement detection, telemetry, or RFID trackers, all of which provide single-channel data streams.^13,14^ More recently, cameras have also been used for recording behavior, supported by advanced machine learning techniques to allow extraction of complex behaviors across conditions using either marker-based or markerless pose estimation.^15,16^

The use of video-monitoring with behavioral classification in home-cage environments can reveal complex rodent behavior longitudinally. Laboratory mice are typically housed in standardized individually ventilated cages (IVCs)^14^ to maximize caging density in animal facilities. These compact cages with low, opaque ceilings make it more challenging to record video with off-the-shelf cameras. Several groups have therefore developed cage designs for monitoring mouse behavior outside of a ventilated rack, using dedicated home-cages to facilitate behavioral monitoring.^17,18^ Such setups often require more space than a traditional IVC home-cage, reducing overall animal housing capacity and requiring specialized care.^19^ Bespoke setups may also introduce new costs or training for institutional animal care and husbandry staff.^12,20,21^

Given the efficiency of IVC housing, efforts have continued to incorporate cameras alongside standard home-cages. One option is to mount a camera above the cage, but opaque lids and food or water apparatuses often obstruct external views.^22^ Furthermore, the clearance for devices mounted externally is very limited (<5 mm) in most commercial racks. One recent solution improved the field of view by using two cage spaces to gain extra height and improve the field of view, but at the expense of reducing housing density by 50%.^11^ Other attempts to place cameras inside the home-cage have employed multiple cage wall inserts to account for reduced visibility due to metal gratings for food and water.^23^

With the goal of effectively monitoring how complex behaviors like nesting evolve as mice age and how transgenic alterations like amyloid pathology alter these trajectories, we developed a home-cage video monitoring system: ***Camus*** (Camera for home-cage recordings of Mus musculus mice). Camus was designed to provide a cost-effective and adaptable pipeline to parse multiple behaviors automatically in high-density IVC cages, allowing for continuous and longitudinal studies. Camus integrates 3D-printed components and commercially available hardware, minimizing modifications to standard IVC housing to decrease the need for additional space. The system can support up to 20 cameras on a single desktop computer, providing infrared-enabled, top-down video perspectives of mouse behavior that are compatible with contemporary behavioral analysis tools. The Camus system takes into account recommendations provided in a recent systematic review on Home-cage monitoring systems to advance the mission of continuous, longitudinal, non-invasive monitoring of an increasing range of animal behavior.^14^

## Results

### Camus home-cage monitoring system

The Camus home-cage monitoring system, detailed in the Methods and in a lab GitHub repository (https://github.com/KimchiLab/Camus), supports video monitoring of laboratory mice without destructive cage modifications. For hardware, Camus combines off-the-shelf cameras and LED lighting with custom designed 3D-printed parts within standard IVC cages (Figure 1a-b). The camera is mounted internally to the ceiling, providing a top-down view of the entire cage floor to capture a wide variety of home-cage behaviors. An unobstructed view of mouse behavior is facilitated by mounting custom food and water holders to the side of the cage (Figure 1b-c). The system as designed supports up to 20 simultaneous cage recordings on one acquisition computer (Figure 1d), at an incremental cost of $250 per cage. Supplementary Video 1 shows a sample Camus recording and Supplementary Video 2 shows simultaneous video streams from 20 cameras.

**Figure 1:**
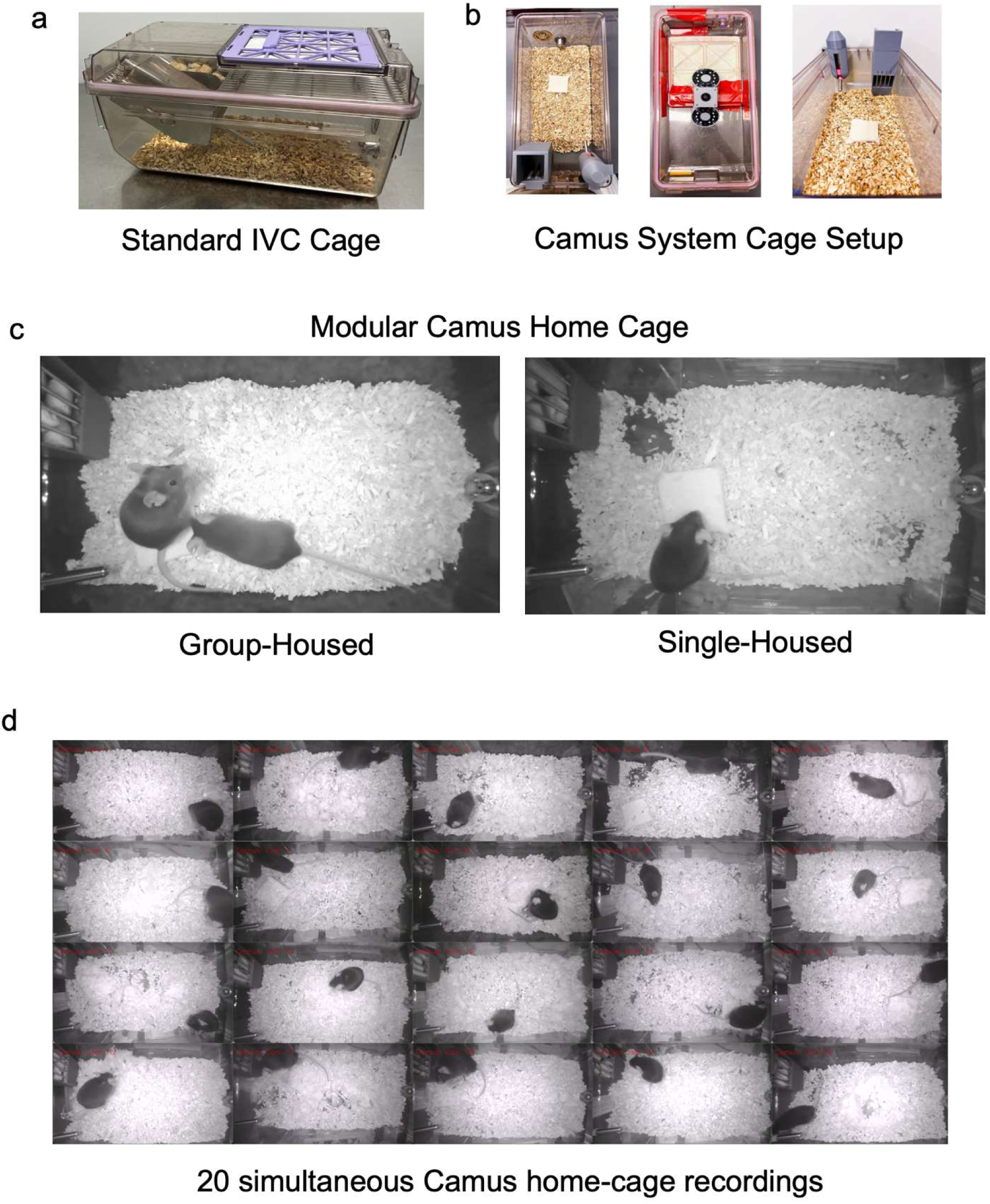
Camus Cage Hardware and Modular Design. **a.** A standard Individually Ventilated Cage (IVC) into which Camus devices are placed. **b.** Different perspective views of the modified Camus home-cage, showing top-down spatial layout of the cag floor (left), bottom-up view of the cage top highlighting the placement of the wide-angle camera with IR LEDs within the custom 3D-printed holder (center), and a deep view illustrating the food hopper, water bottle, and nestlet (right). **c.** Video snapshots depicting pair-housed (left) and single-housed (right) mice in Camus home-cages. **d.** Screenshot showing simultaneous recordings from 20 individual Camus home-cages on a single computer.

### Evaluation of mouse weights in the Camus environment

We first assessed whether mice maintained similar weights in the Camus environment with its modified feeding and drinking devices, compared to standard home-cages. We monitored weights daily for three days after placing mice into new cages of each type (Figure 2a-b). Although female mice weighed less than male mice (linear mixed-effects model (LMEM), significant main of Sex, F_1,12_ = 25.06, p = 0.0003), weights remained stable for both sexes across multiple days (Figure 2c). There was no significant main of Cage-Environment on weight (F_1,111_ = 1.34, p = 0.249), nor an interaction between Days x Cage-Environment (F_1,111_ = 0.18, p = 0.674), demonstrating weight stability in the Camus environment.

**Figure 2:**
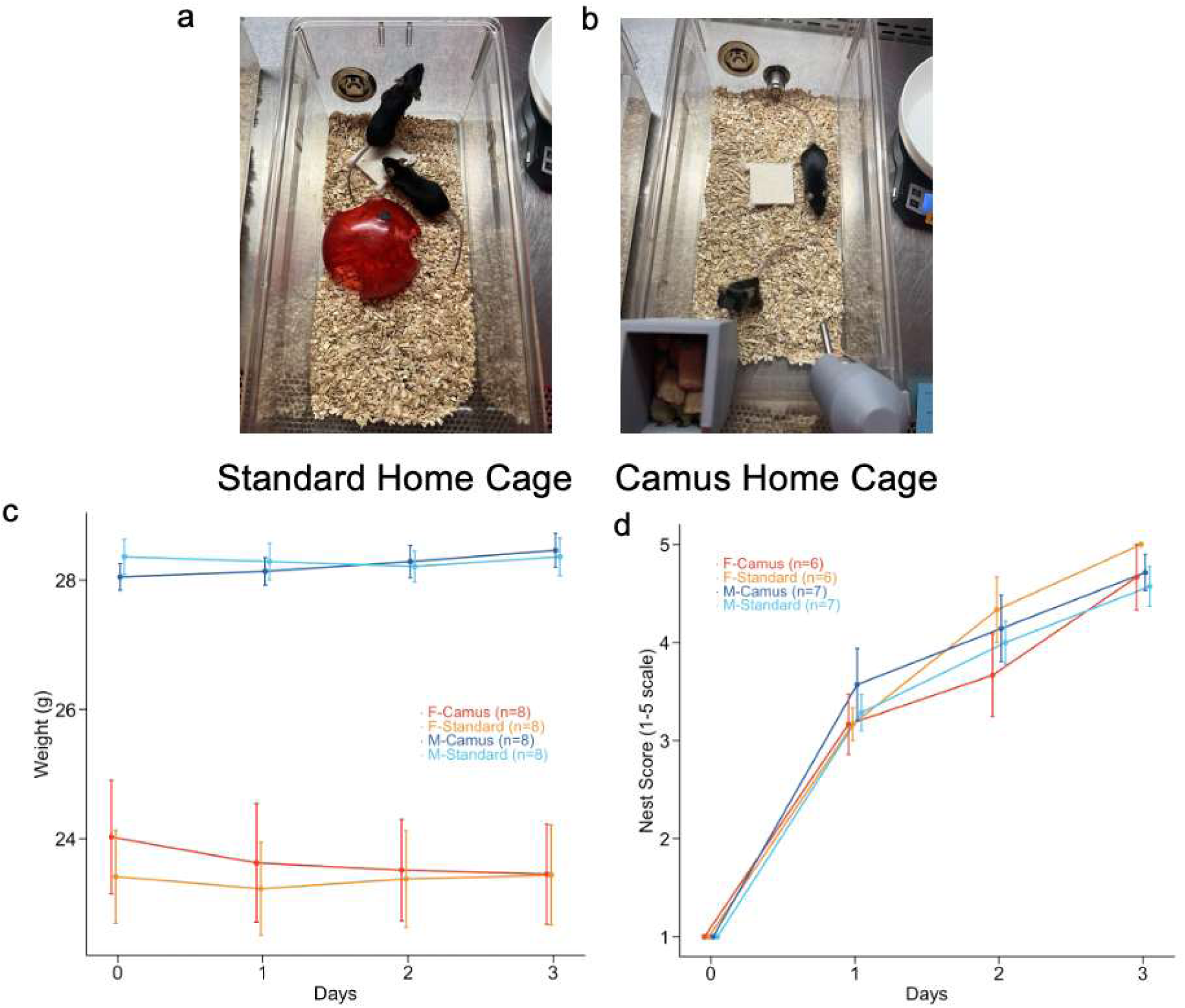
Validating weight and nest building in Camus Cage compared to traditional IVC Home-Cage. **a.** Top-down view of standard home-cage with two mice. Food and water were provided in a wire grating lid, which has been removed for visibility for this photo. **b.** Top-down view of Camus home-cage with two mice. Food and water were provided in the 3D printed food hopper (left) and via an internally mounted sipper tube water bottle (right). **c.** Mouse weights were recorded daily across three days after placement in Camus or standard home-cages (n = 16 mice; within-subject design, counter-balanced order). Data were analyzed using a linear mixed-effects model (LMEM) with fixed effects of Day, Cage-Environment, and Sex, and a random intercept for Subject. Although females weighed significantly less than males (F_1,12_ = 25.06, p = 0.0003), weight remained stable across Days (F_1,111_ = 0.044, p = 0.834) and did not differ between Camus and standard housing (F_1,111_ = 1.34, p = 0.249). There was no interaction between Day and Cage-Environment (F_1,111_ = 0.18, p = 0.674). **d.** Nest-building was scored daily on a 1-5 scale for pairs of mice in both cage environments (n = 13 cohorts; within-subject design, counter-balanced cage-environment order). Data were analyzed using a linear mixed-effects model (LMEM) with fixed effects of Day, Cage-Environment, and Sex, and a random intercept for Subjects. There were no significant main effects of Cage-Environment (F_1,60 =_ 0.129, p = 0.720) or Sex (F_1,11_ = 0.043, p = 0.8395). There was no significant interaction between Cage-Environment × Day (F_2,60_ = 0.614, p = 0.544). These results indicate that nesting ability was similar between the two environments over days, supporting the Camus system’s preservation of normal nesting behavior.

### Evaluation of mouse nesting in the Camus environment

We next determined whether mice displayed similar nest-building behavior in the Camus environment as in standard home-cages. We place mice into a new cage-environment of each type with a compressed cotton nestlet. We monitored and scored nests daily for three days, using a modified nest scale where 1 indicates no nest and 5 indicates a complete nest (Figure 2d).^24^ Linear mixed-effects model analysis revealed no significant effects on nest building of Cage-Environment (F_1,60_ = 0.129, p = 0.720), Sex (F_1,11_ = 0.043, p = 0.8395), or Cage-Environment × Days interaction (F_2,60_ = 0.614, p = 0.544), indicating similar nest building between the standard and Camus environments over days.

### Camus video recordings

We have used the Camus system to record 1,198 multi-day sessions from 336 unique mice across multiple experiments. Recordings were performed using Python 3.11.12 scripts, using OpenCV-Python 4.6.0 on a Windows desktop computer. Video data was recorded with a resolution of 640×360 pixels at a frame rate of 30 frames per second, with file sizes for one-hour recordings having a mean and standard deviation of 0.861 ± 0.097 GB/hour (calculated from 5,757 total AVI files). We captured ∼99% of expected frames throughout all recording sessions, ensuring minimal data loss for downstream analyses.

### Camus keypoint estimation

Videos were analyzed using keypoint based tracking using a trained DeepLabCut computer vision model, developed using manual labeling of 2,016 frames taken from 213 videos representing 44 different mice. 36 labeled keypoints included 22 mouse keypoints and 14 environmental markers, such as the food hopper, waterspout, cage ventilation port, and nesting material. All keypoints are presented in Supplementary Figure 1. After 950,000 iterations, the training error with likelihood cutoff of p>0.6 was 2.95 for training images, and 9.33 for test images (Supplementary Table 1).

Body keypoints from a single mouse alongside nesting and other environmental keypoints are depicted over time in Figure 4. Supplementary Video 3 shows a mouse with an overlaid skeleton of extracted keypoints, demonstrating the outputs of the pose estimation model. This DeepLabCut model is available in the Camus repository.

### Camus records circadian rhythms in locomotion, feeding, and drinking, and reveals sex-dependent activity

We used extracted keypoints to study the circadian activity of mice housed in the Camus system. Mice showed robust circadian organization of typical behaviors, consistent with normal nocturnal physiology (Figure 3).

**Figure 3:**
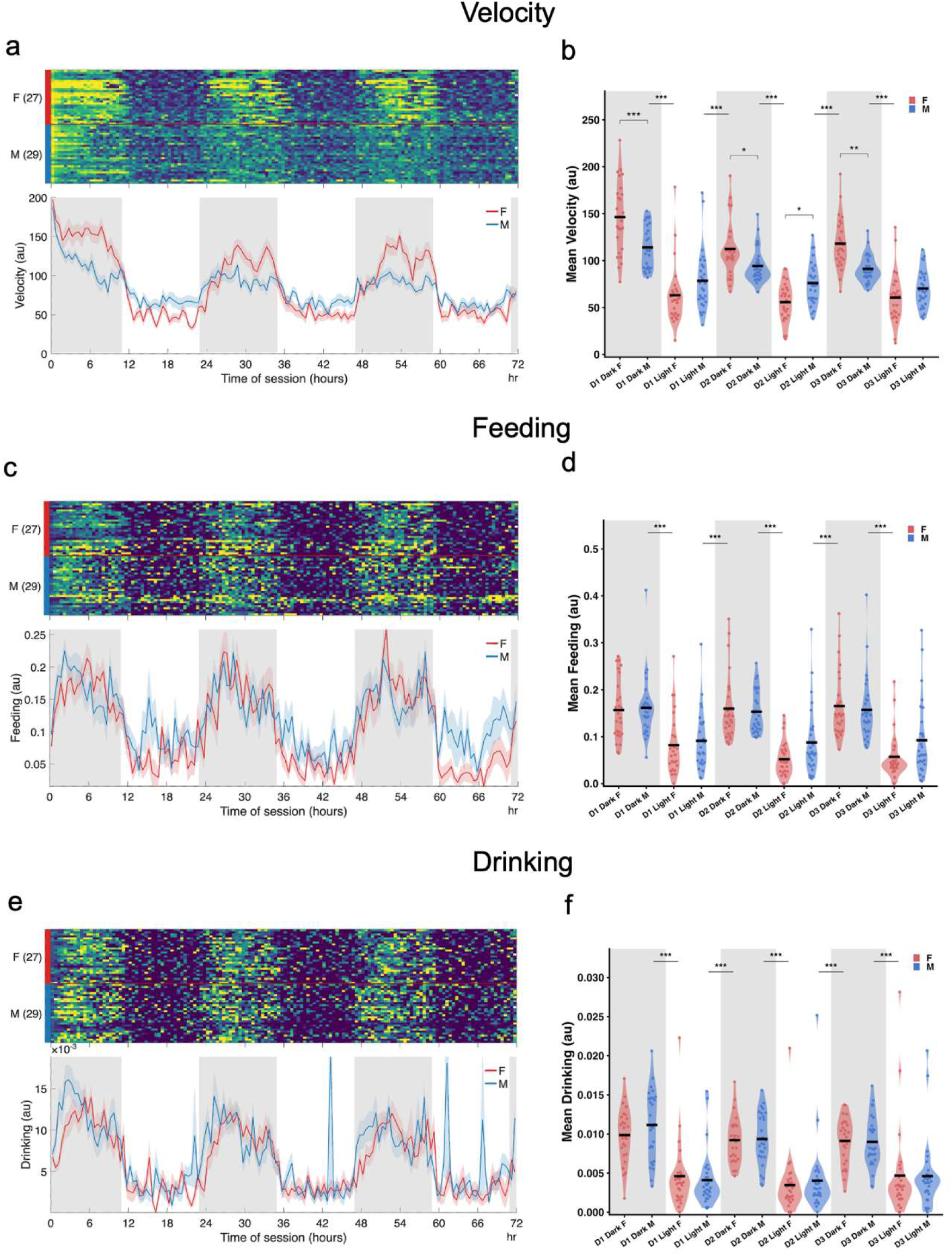
Sex differences in Camus home-cage behavior across the circadian cycle. **a-b. Locomotion Velocity. a.** The heatmap depicts movement of individual mice within each row, summarized as a series of 30-minute bins over 72 hours. Brighter colors indicated increased velocity. Female mice are above (F, red bar), and male mice are below (M, blue bar). The lower plot shows average velocity by sex (lines = means, shading = standard error). **b.** Locomotion velocity differed significantly across 12 hour Windows (LMEM, p <0.0001). At each transition across successive windows, mice moved more during dark windows compared to light windows (Sidak-adjusted post-hoc tests of 5 transitions, all p<0.0001, represented as horizontal line across dark/light transitions). Females moved more than males during all dark windows (LMEM, Sex × Window, p<0.0001; Holm-adjusted post-hoc tests, all p<0.05, represented as overlying comparison bracket within each window). Females also moved less than males during the light window on Day 2 (D2 Light F vs. M, Holm-adjusted post-hoc tests, p = 0.0256). Significance code for post-hoc tests: *** <0.001, ** <0.01, * <0.05. **c-d. Feeding. c.** The heatmap depicts feeding bouts defined as closeness to the food hopper over 72 hours, with conventions as in panel a. **d.** Feeding differed significantly across Windows (LMEM, p <0.0001). At each transition across the six successive windows, mice interacted with the food hopper more during dark windows compared to light windows (emmeans contrasts Sidak-adjusted comparison of 5 transitions, all p<0.0001). There was no significant effect of Sex nor interaction between Sex × Window. Conventions as in panel b. **e-f. Drinking. e.** The heatmap depicts drinking bouts defined as closeness to the water spout over 72 hours, with conventions as in panel a. **f.** Drinking differed significantly across Windows (LMEM, p <0.0001). Post-hoc tests showed that at each transition across the six successive windows, mice interacted with the water spout more during dark windows compared to light windows (emmeans contrasts Sidak-adjusted comparison of 5 transitions, all p<0.0001). Conventions as in panel b.

Mouse locomotion, derived from the movement of a centralized mouse body keypoint, differed across Windows after placement in Camus (LMEM, F_5,280.38_ =87.624, p <0.0001) (Figure 3a-b). Post-hoc tests showed that at each transition across the six successive windows, mice moved more during dark windows compared to light windows (emmeans contrasts Sidak-adjusted comparison of 5 transitions, |t_280_|= 8.082-14.736, all p<0.0001). Although female and male mice overall moved at similar amounts (no significant LMEM effect of sex on locomotion: F_1,38.8_ = 1.033, p = 0.3158), there was an interaction between Sex × Window on locomotion (LMEM, F_5,280.38_ = 16.387, p <0.0001). Post-hoc contrasts revealed that females moved more than males during all dark windows (emmeans contrasts Holm-adjusted, t_147.29_ = 2.453-4.394, all p<0.05). Females also moved less than males during the light window on Day 2 (emmeans contrasts Holm-adjusted, t_147.29_ =−2.767, p = 0.0256).

Feeding behavior, measured by posture relative to the food hopper, also differed across Windows (LMEM, F_5,280.38_ =44.71, p <0.0001) (Figure 3c-d). Post-hoc tests showed that at each transition across the six successive windows, mice interacted with the food hopper more during dark windows compared to light windows (emmeans contrasts Sidak-adjusted comparison of 5 transitions, |t_280_|= 7.292-9.574, all p<0.0001). We did not observe a significant effect of Sex (LMEM, F_1,33.06_ = 0.849, p = 0.3634), nor an interaction between Sex × Window (LMEM, F_5,280.38_ = 2.108, p = 0.0646).

Drinking behavior, measured by posture relative to the water spout, also differed across Windows (LMEM, F_5,280.38_ = 43.350, p<0.0001) (Figure 3e-f). Post-hoc tests showed that at each transition across the six successive windows, mice interacted with the water spout more during dark windows compared to light windows (emmeans contrasts Sidak-adjusted comparison of 5 transitions, |t_280_|= 6.848-9.577, all p<0.0001). We did not observe a significant effect of Sex (LMEM, F_1,42.66_ = 0.029, p = 0.8667), nor an interaction between Sex × Window (LMEM, F_5,280.38_ = 0.489, p = 0.7844). Together, these findings show that Camus captures expected circadian patterns in locomotion, feeding, and drinking behaviors, while also detecting sex-specific differences in locomotion that vary between light and dark circadian windows.

### Camus reveals nest-building dynamics and sex differences in nest construction

We used Camus to study nest-building dynamics within the home-cage, by tracking the disappearance of the compressed nestlet and the emergence of a round, crater-like nest (Figure 4a-b). To study nesting quantitatively, nestlet disappearance was measured using the Nestlet keypoint likelihood from the DeepLabCut model, and nest appearance was measured using the mean likelihood of all eight-cardinal direction Nest keypoints. We first validated these quantitative metrics of nesting behavior by comparing DeepLabCut based nesting metrics (Nestlet likelihood and Nest likelihood) with manually rated consensus nest scores (Supplementary Figure 3). The initial interrater reliability for visual scoring of nests from video frames was 0.851 (Krippendorff’s alpha), and consensus was subsequently achieved for all frames. The average Nestlet likelihood was lower at higher nest scores, and the average Nest likelihood monotonically increased with higher nest scores (Supplementary Figure 3).

**Figure 4:**
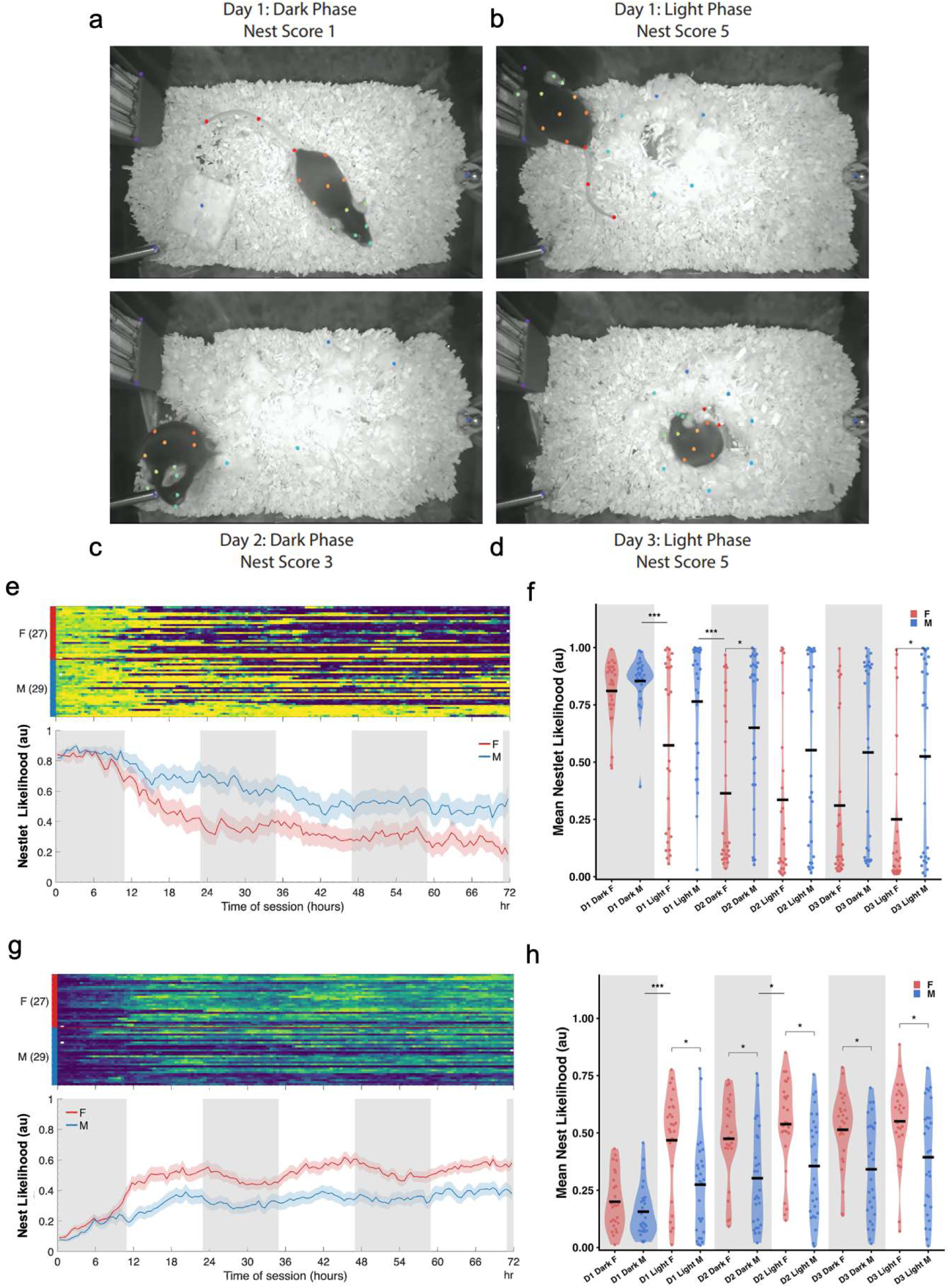
Sex-differences in Nest-building dynamics across the circadian cycle. **a-d.** Representative frames illustrate keypoint estimation for mouse tracking and nest dynamics in Camus videos of the same mouse across multiple days, using DeepLabCut. **a.** Day 1, Dark Phase/Window: points were identified for the mouse and the nestlet; this image corresponds to a nest score of 1. **b.** Day 1, Light Phase/Window: example of a frame corresponding to a well-formed, crater-like nest structure (score of 5). **c.** Day 2, Dark Phase/Window: example of a frame corresponding to a partially disrupted nest (score of 3), reflecting nest displacement during active dark periods. **d.** Day 3, Light Phase/Window: example of a frame corresponding to a restored, high-quality nest (score of 5). **e-f. Nestlet likelihood.** The heatmap shows per-session Nestlet likelihood in 30-minute bins over 72 h. Female mice (F, red bar) are above, and male mice are below (M, blue bar). The lower plot shows average velocity by sex (lines = means, shading = standard error). **f.** Nestlet likelihood differed significantly across Windows (LMEM, p<0.0001). The first two window transitions showed significant decreases in nestlet likelihood (emmeans contrasts Sidak-adjusted, Day 1 Dark to Day 1 Light, p = 0.0001; Day 1 Light to Day 2 Dark, p = 0.0001, represented as horizontal line across dark/light transitions). There was also a main effect of Sex (LMEM,p = 0.0151) on Nestlet and a significant Sex × Window interaction (LMEM, p = 0.0227). During two isolated windows, females had lower Nestlet likelihoods than males (emmeans contrasts Holm-adjusted, Day 2 Dark, p = 0.0217; Day 3 Light, p = 0.0252 represented as overlying comparison bracket within each window). Significance code for post-hoc tests: *** <0.001, ** <0.01, * <0.05. **g-h. Nest likelihood.** The heatmap shows per-session Nest likelihood in 30-minute bins over 72 hour (conventions as in e). **h.** Nest likelihood differed significantly with Windows (LMEM, p<0.0001). The first two dark to light transitions showed significant increases in Nest likelihood (emmeans contrasts Sidak-adjusted, Day 1 Dark to Light, p < 0.0001; Day 2 Dark to Light, p = 0.0356). There was also a significant effect of Sex (LMEM, p = 0.0087), as well as a significant Sex × Window interaction (LMEM, p<0.0001). Except for the first window, Nest building in females exceeded males in all five subsequent windows, emmeans contrasts holm-adjusted comparison of F-M in 5 windows, all p<0.05. Conventions as in f.

We then tracked quantitative nesting metrics over time. Nestlet likelihood decreased over time after placement in Camus cages (Figure 4e-f), as reflected by a significant effect of Window on Nestlet likelihood (LMEM, F_5,280.38_ = 41.087, p<0.0001). Post-hoc tests were done across six successive windows showing each dark-light transition. The first two transitions showed significant decreases in nestlet likelihood (emmeans contrasts Sidak-adjusted, Day 1 Dark to Light: t_280_ =−4.298, p = 0.0001; Day 1 Light to Day 2 Dark: t_280_ =-4.261, p = 0.0001), indicating that nestlet shredding primarily occurred in the first 24-36 hours. There was a main effect of Sex (LMEM, F_1,34.97_ = 6.536, p = 0.0151) on Nestlet likelihood, with increased use of the nestlet by female mice. There was also a significant Sex × Window interaction (LMEM, F_5,280.38_ = 2.663, p = 0.0227). During two isolated windows, females had lower Nestlet likelihoods than males, suggesting time-specific increased nestlet use by females (emmeans contrasts Holm-adjusted, Day 2 Dark: t_67.12_ =−3.016, p = 0.0217; Day 3 Light: t_67.12_ =−2.900, p = 0.0252; all other windows were not significantly different: p > 0.05).

Complementary to the decreasing Nestlet likelihood, we saw an increasing Nest likelihood over time (Figure 4g-h), with a significant effect of Window on Nest likelihood (LMEM, F_5,280.38_ = 48.294, p<0.0001). The first two dark to light transitions showed significant increases in Nest likelihood (emmeans contrasts Sidak-adjusted, Day 1 Dark to Light: t_280_ = 8.948, p < 0.0001; Day 2 Dark to Light: t_280_ 2.706, p = 0.0356), indicating early building of nests. There was also a significant effect of Sex (LMEM, F_1,42.83_ = 7.557, p = 0.0087), with increased nest building by females. And there was a significant Sex × Window interaction (LMEM, F_5,305_ = 5.703, p<0.0001). Post-hoc tests showed that after the first Day 1 Dark window, Female mice had greater nest likelihood in all subsequent windows than Male mice (emmeans contrasts holm-adjusted comparison of F-M in 5 windows, t_70.93_= 2.476-3.105, all p<0.05).

We sometimes observed degradation and then reconstruction of nests over subsequent days (Figure 4c-d), including for manually rated Nest scores across 4 mice (Supplementary Figure 4). To characterize nesting dynamics over time, we compared nests in 6-hour windows (Supplementary Figure 5). A model with all twelve 6-hour Windows also showed a significant effect of Sex × Window interaction (LMEM, F_11,580.88_ = 3.341, p=0.0002). Post-hoc testing on the final 6 hours of each circadian phase revealed that Female mice consistently alternated Nest likelihoods, with Nest likelihood rising following each dark-to-light transition (emmeans contrasts holm-adjusted, Day 1, t_581.73_ = 7.807, p < 0.0001; Day 2, t_581.73_ = 3.718, p =0.0009; Day 3, t_581.73_ = 2.346, p = 0.0386) and falling following each light-to-dark transition (emmeans contrasts holm-adjusted, Day 2, t_581.73_ = −2.086, p = 0.0386; Day 3, t_581.73_ = −2.536, p = 0.0344). Male mice showed only an initial rise in Nest likelihood following the first dark-to-light transition (emmeans contrasts holm-adjusted, Day 1, t_581.89_ = 4.842, p < 0.0001), with no subsequent window transitions reaching significance (emmeans contrasts holm-adjusted, |t_581.89_|= 0.276-1.704, all p ≥ 0.3554). Together, these findings show that nest-building provides a continued measure of organized home-cage behavior that differs between male and female mice.

### Camus reveals a selective, age-progressive nesting deficit in 5xFAD mice

We finally used Camus to monitor activity in sex-balanced, transgenic 5xFAD Alzheimer amyloid mice and wild-type (WT) littermates across three age ranges: 2-6, 6-12, and 12-18 months. For aging analyses, each session was summarized as a mean value from the full 72-hour recording. Linear mixed effects models were performed on these mean values using Age as a main effect and Mouse as a random effect. WT and 5xFAD mice were modeled separately. For both 5xFAD and WT mice, locomotor activity remained stable across ages. There was no effect of Age on mean Locomotion for both WT (LMEM, F_2,43.7_ = 0.621, p = 0.5420; Figure 5a&c) and 5xFAD mice (LMEM, F_2,43.35_ = 0.867, p = 0.4275; Figure 5b&d), suggesting a stable motor baseline for both WT and 5xFAD mice prior to examination of more detailed behavior, such as nest building.

**Figure 5:**
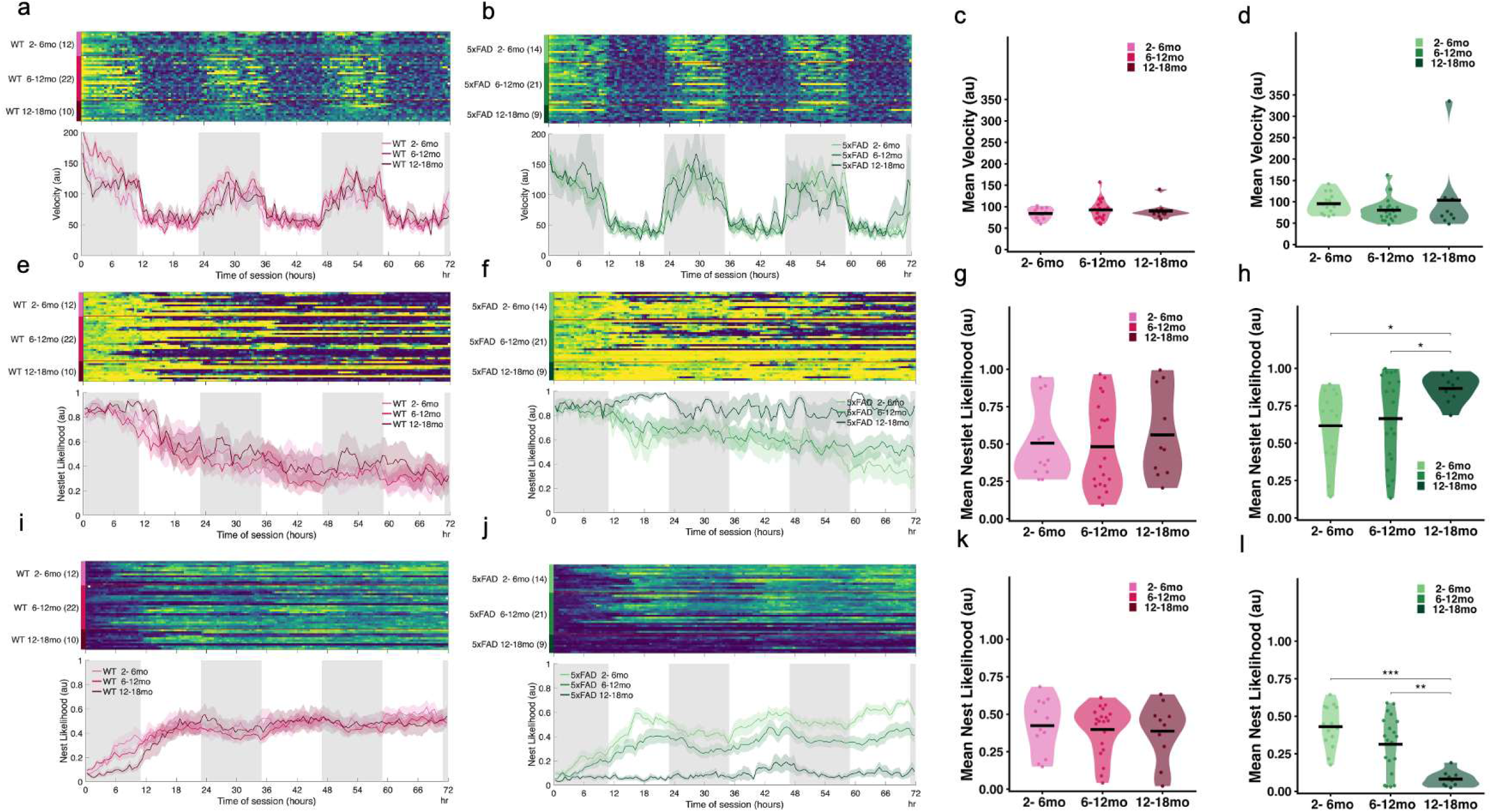
Behavioral aging in WT versus transgenic 5xFAD mice. **a-d. Locomotor Velocity.** The heatmaps in **a** (WT) and **b** (5xFAD) depict individual locomotor velocity (30-minute bins over 72 hours), with brighter colors indicating increased velocity. The lower panel shows average group velocity (lines = means, shading = standard error). WT (pink) and 5xFAD (green) mice were recorded across 2-6-, 6-12-, and 12-18-month age groups. Mean velocity over the entire session did not change with age in either group (**c**=WT, **d**=5xFAD). Each dot in the violin plot represents mean velocity over the 72-Hour recording window. Significance code for post-hoc tests: *** <0.001, ** <0.01, * <0.05 are represented with overlying comparison brackets. **e-h. Nestlet likelihood.** The heatmaps in **e** and **f** show per-session Nestlet likelihoods (30-minute bins over 72 h), with brighter colors indicating increased Nestlet likelihood (conventions as in a, b). **g.** Nestlet likelihood did not change with age in WT mice (LMEM, p = 0.7916). **h.** 5xFAD mice showed a difference in mean Nestlet likelihood by Age (LMEM, p = 0.0138). Post-hoc tests revealed a significant increase in remaining Nestlet likelihood between for the older, 12-18-month mice compared to younger groups (emmeans contrast Tukey-adjusted, both p<0.05). Conventions as in c-d. **i-l. Nest likelihood.** The heatmaps in **i** and **j** show per-session Nest likelihood (30-minute bins over 72 h), with brighter colors indicating increased Nest likelihood (conventions as in a, b). **k.** Nest likelihood did not change with age in WT mice (LMEM, p = 0.8567). **l.** 5xFAD mice showed decreased nest building with age (LMEM, p < 0.0001). Post-hoc tests revealed that 12-18-month-old 5xFAD mice were significantly less likely to build Nests compared to younger groups (emmeans contrast Tukey-adjusted, both p<0.01). Conventions as in c-d.

In WT mice, nesting behavior remained stable across these ages, both when analyzing Nestlet likelihood (LMEM no effect of Age, F_2,43.7_ = 0.235, p = 0.7916) (Figure 5e&g) and Nest likelihood (LMEM no effect of Age, F_2,40.71_= 0.155, p = 0.8567) (Figure 5i&k). In contrast, in 5xFAD mice, nest-building declined with age. There was a significant effect of Age on Nestlet likelihood in 5xFAD mice (LMEM, F_2,29.71_ = 4.969, p = 0.0138). Post-hoc tests revealed that 12-18-month-old 5xFAD mice were significantly less likely to use up the Nestlet than 2-6-month-old (emmeans contrast Tukey-adjusted, t_29.7_ =−2.960, p=0.0160) and 6-12-month-old mice (emmeans contrast Tukey-adjusted, t_38.2_=−2.658, p=0.0301). Nest likelihood also showed a significant effect of Age in 5xFAD mice (LMEM, F_2,37.48_ = 13.683, p < 0.0001; Figure 5j&l). Post-hoc tests revealed that 12-18-month-old 5xFAD mice were significantly less likely to build their Nests than 2-6-month-old (emmeans contrast Tukey-adjusted, t_37.5_ = 5.221, p<0.0001) and 6-12-month-old mice (emmeans contrast Tukey-adjusted, t_44.0_= 3.632, p=0.0021). Thus, older 5xFAD mice showed a loss of organized nest-building despite preserved locomotor activity, whereas age-matched WT mice did not. These findings identify a genotype-specific, age-dependent behavioral deficit that is more apparent in nest-building than in overall movement.

## Discussion

We developed Camus as an open-source platform for continuous video monitoring of spontaneous mouse behavior in standard individually ventilated cages. The system combines low-cost cameras, infrared illumination, adaptable 3D-printed components, and computer-vision analysis while maintaining the general organization and density of conventional laboratory housing. A single desktop computer recorded up to 20 cages in parallel at 30 frames per second and captured more than 99% of expected frames. Under the conditions tested, mice housed in Camus cages did not differ from mice in conventional cages in body weight or nest quality. These findings support the feasibility of sustained, parallel behavioral monitoring without specialized chambers or major changes to routine husbandry.

A central goal of Camus was to make continuous behavioral measurements accessible within the cages in which laboratory mice already live. Mounting the camera and infrared illumination within the cage lid avoided external camera setups, specialized racks, and the additional space required by existing systems. The use of commercially available cameras, modifiable 3D-printed components, and open-source acquisition and analysis software also supported a modular design. This allows individual parts of the system such as 3D printed food hoppers to be replaced or adapted as experimental needs change, and can lead to inclusion of other open-sourced 3D printed devices such as FED3 to measure food intake in the home-cage more precisely.^25^ The IR LEDs emit light at 850nm at an intensity which allows for effective monitoring without compromising the darkness needed for the active window.^26^ Camus is currently designed for a specific IVC product, but its modular design should allow adaptation to other cages, cameras, illumination sources, and behavioral apparatuses. Any such modification might require additional validation because changes in viewing angle, illumination, or cage structure might affect both animal behavior and computer-vision performance.

Continuous video is most informative when paired with analytical methods that preserve the temporal organization of behavior.^27,28^ Marker-less keypoint-estimation using DeepLabCut^29^ model tracked both the mouse and relevant landmarks within the cage, allowing analysis of locomotion, feeding, drinking, and nest building to be measured from the same recordings, with potential of expanding to the detection of additional features such as grooming. Our system allows for the collection of multi-animal videos. However, given the challenges and ongoing development of multi-animal tracking in software such as DeepLabCut and SLEAP, which sometimes requires manual review that would be challenging in large-scale recordings, our current pose-estimation model was focused on single-housed mice at this time.^29,30^

Across the typical single-housed 72-hour recordings, Camus demonstrated the expected nocturnal patterns in locomotion, feeding, and drinking.^31,32^ The system also detected an interaction between sex and circadian window: female mice were more active than male mice during the dark window but less active during some light windows, consistent with previously reported sex differences in mouse activity.^33,34^ Because these opposing differences were lost when activity was averaged across the full light-dark cycle, this finding illustrates an important advantage of continuous monitoring. Behavioral differences that may depend on circadian window or behavioral state can be obscured when measurements are pooled across time.

Nest construction provided a particularly clear example of the information gained by treating behavior as a dynamic process rather than a single endpoint. Conventional nest-building tests often summarize behavior into one score collected after an overnight interval.^35^ In contrast, Camus tracked the disappearance of the intact nestlet and the formation, disruption, and reconstruction of nests throughout the recording, especially for female mice. Nest likelihood increased during the light window, when nests were most cohesive, and was generally higher in females after the first dark window.^36,37^ For females, we observed cycles of construction and disruption which was most notable in the second halves of the circadian windows. These findings demonstrate that at least in female mice, a nest is not simply a static final product but the result of behavior that continues across successive light and dark periods.^36^ Continuous monitoring can reveal when construction begins, how quickly nesting material is incorporated, whether the structure remains cohesive, and how it changes across the circadian cycle. Such features could not be recovered from a single terminal score.

Applying Camus across age-groups in the 5xFAD mouse model showed how temporal and behavioral detail may expand disease-model phenotyping.^38–41^ Locomotor activity did not decline significantly with age in either 5xFAD or wild-type mice during the studied ages, and nest-related measures remained stable in wild-type littermates.^40,42^ In 5xFAD mice, however, nesting declined with age. This may be driven by the progressive age-related amyloid plaque accumulation in 5xFAD mouse cortex and hippocampus and age-related microgliosis.^38^ The lack of a corresponding decline in locomotion makes a simple explanation based on generalized motor impairment less likely.

Nest construction should nevertheless be interpreted as an integrated behavioral phenotype rather than a specific measure of a single cognitive process. Successful nest-building requires mice to engage with the material, shred and transport it, organize it spatially, and continue modifying the structure over time. Performance may therefore be influenced by motivation, thermoregulation, somato-sensation, motor function, arousal, and the organization of behavioral sequences ^43,44^ This sensitivity to multiple neural and physiological processes makes nesting useful for detecting altered function but may limit its mechanistic specificity. Future studies combining Camus’s measurements with targeted perturbations, physiological recordings, and complementary behavioral assays will be needed to determine which components of nest-building are altered by aging and amyloid pathology.

Several technical limitations should be considered. The top-down camera provided broad coverage of horizontal movement and interactions with food, water, and nesting material but may not fully capture vertical behaviors such as rearing or climbing, and inferior postures such as detailed gait analysis. Feeding and drinking were inferred from the proximity of tracked facial keypoints to the corresponding apparatus. These measures therefore estimate food- and water-directed behavior rather than directly confirming consumption. Similarly, Nestlet presence and Nest likelihood are computer-vision-derived measures of material processing and nest formation rather than direct measurements of every construction movement. Performance may vary across cage configurations, bedding conditions, early development, and other mouse coat colors. External validation and transparent reporting of model performance will therefore be important for broader adoption. Continuous recording also generates large volumes of video data, requiring users to balance temporal resolution against storage and computational demands.

When desired, storage of raw videos allows for continued analysis including new behavioral features or the use of alternate pose estimation techniques. But storage of raw video may create significant data storage requirements. At 0.861 GB/hr/cage, 20 cages generate 413 GB per 24 hours of recording, which can require large storage requirements for longer high-throughput longitudinal recordings. Data storage could be optimized with lower spatial or temporal resolution videos, higher compression algorithms, or by solely retaining keypoints or extracted features following initial processing. Keypoint based data can still facilitate future work using sequence-based classifiers and unsupervised methods to identify additional behavioral patterns and state transitions from keypoints.^45^

In summary, Camus enables continuous and parallel monitoring of spontaneous mouse behavior in standard laboratory housing. Its main advantage is scale with temporal resolution: behaviors that are typically measured as isolated endpoints can be studied as structured processes unfolding across hours, days, circadian windows, sex, age, and disease state. By facilitating raw video recording in the home cage and providing adaptable hardware designs, open-source software, and extensible analysis tools, Camus provides a scalable foundation for continuous behavioral phenotyping that can evolve with new experimental questions and computational methods.

## Materials and Methods

### Animals

All presented behavioral data were collected from Alzheimer’s amyloid model 5xFAD and wild-type littermates. Hemizygous transgenic 5xFAD male mice on a congenic C57BL/6J background (courtesy of Dr. Robert Vassar), were bred with female, wild-type C57BL/6J mice (Jax 000664).^46^ Pups were weaned and housed by sex at 21 days. Mice were genotyped for 5xFAD transgenic markers (Transnetyx) and described as “5xFAD” or “wild-type” (WT) depending on results. Wild-type mice were 2-24 months old at the time of video recordings. Transgenic 5xFAD mice were 2-18 months old at the time of video recordings and compared to age-matched wild-type mice for joint analyses. Sample size was determined by use of mice in ongoing experiments and are presented here as a sample of convenience.

We additionally recorded videos from an Alzheimer’s tauopathy model TauP301S (Jax 008169) mice on a B6C3F1/J background (Jackson Labs). Mice were bred as one transgenic male paired with two female, wild-type C57BL/6J mice (Jax 000664). Pups were weaned and housed by sex at 21 days and displayed either black or agouti coat colors. These videos were only used to train the keypoint estimation DeepLabCut model described below and are not presented in behavioral analyses.

### Standard Animal Facility Housing

Mice were housed in Allentown NexGen IVC home-cages. A reverse 12/12 daylight cycle was used, with lights off from 9am to 9pm and red lights used when working during these hours. The rodent housing facility was maintained at a mean temperature of 72 ± 2 °F and a relative humidity range of 30% to 70%. Mice were housed within university animal areas, and all procedures were approved by the Institutional Animal Care and Use Committee (IACUC). Cages contained compressed nestlet material (compacted cotton fiber 5x5 cm square, Ancare) and bedding (Aspen Sani Chips), and mice were fed standard laboratory diet (Teklad LM-485 Mouse sterilizable diet).

### Camus Home-cage Design

20 mouse cages, housing 1-2 mice, were all connected to a single desktop computer for simultaneous camera recordings. The wire grating cage top that would typically hold laboratory chow and water bottles was removed because the grating would visually obstruct the camera view. Food was instead provided using a 3D printed food hopper containing the standard laboratory diet held by thin metal rods (Grainger Stainless Steel Rod: 303, 3/32” outer diameter) for robustness against gnawing. Water was provided from a 3D printed water bottle holder for a commercial water bottle (80 mL, roll ball sipper, purchased on AliExpress) designed for domestic pets. At a typical water intake of 8 ml/30 g body weight for C57BL/6J mice, our water bottles allow for a continuous weeklong recording without a bottle change for single mice.^47^ 3D printed parts were designed in Blender 3.5 and are open-source and available via the repository under a CERN-OHL-S v2 open-source license. Parts were printed using Polymaker Gray PLA+ 1.75mm filament on a Prusa MK3S+ printer. The modified Camus home-cages can be further checked for BSL2 work by doing seal validation for barrier integrity

Camus cages were outfitted with a USB camera with a low-distortion, 120° lens (ELP-USBFHD05MT-RH120, Shenzhen Alipu) with two infrared LED panels aligned at 35° angles along the long axis of the cage, in a 3D printed housing using polylactic acid (PLA) on a Prusa MK3S+ printer. The camera was taped (Poly Stucco Tape 3262) to a standard cage top (Allentown Edge Cage System with Micro Barrier Top) with cables routed out of the cage top in a protected manner. The approximate print time for all parts for a single cage was 14 hours using a Prusa MK3S+.

Additional materials such as USB cables and infrared 850nm LED panels were purchased as readily available parts from standard commercial vendors as documented in the repository parts list. The cost per Camus cage setup was $250 each (USD) across 20 cages including the shared cost of the computer to record all videos simultaneously. Rack level power draw from the computer was 75 W for 20 cameras with dual IR LED boards. Assembly time per camera and cage was approximately one hour but could vary based on soldering experience. Although designed to fit an Allentown NexGen IVC mouse cage, 3D design files are editable to adapt to cages of different sizes. The food-hopper, water-bottle holder, and the water bottles are multi-use and are cleaned and sanitized after each session using IACUC-approved standard operating procedure where all parts are submerged in water with soap, rinsed, and then cleaned by spraying with 70% isopropyl alcohol solution.

### Camus Sessions

We started session video recordings between 9:30-10:30am (lights off at 9:00am). Mice were placed in a fresh IVC cage bottom with a new compacted cotton fiber square nestlet for nest building. Fresh food and water were provided in the food hoppers and water bottle holders. The cage bottom was then covered with the Camus cage top and then placed into the IVC rack. Mice remained housed in Camus cages for at least 72 hours and checked by husbandry staff daily.

Recordings do not have to be limited to 72-hours and mice can be monitored continuously with weekly food/water refills and biwkeely cage-bottom changes. Cage tops were sprayed and wiped using a chlorine dioxide disinfectant at least weekly or when mice were removed from cages.

### Measurement of Mouse Weights in Camus Cages

To ensure that mice weights were similar for those housed in Camus compared to standard cages, we placed mice into new Camus or standard cages and recorded daily weights at the same time each day for 3 days (n = 16, 8 males & 8 females). Mice were then moved to a fresh cage of the other type, in counter-balanced fashion. Each mouse served as its within-subject control. Linear mixed effects models were used to analyze the effects of cage environment and sex on weight over days (*lmer* in version R 4.5.3).^48^

### Qualitative Measurement of Nesting in Camus Cages

To ensure that nesting behavior was similar in Camus cages, we placed mice into Camus or standard cages and assessed nest building at the same time each day for 3 days. Nests were scored using a previously described scale of 1-5, where 5 is the most developed nest.^24^ We used a modified version of the scale where we assessed only the nesting structure and did not take weight of nestlet shred into account as described in the previously published scale, so as not to disturb the home-cage environment. 13 pairs (6 Female and 7 Male) of mice were placed in a fresh cage for 3 days, after which they were switched to a fresh cage of the opposite type in counter-balanced fashion. Each mouse pair served as its within-group control. Linear mixed effects models were used to analyze the effects of cage environment and sex on nesting scores over days.

### Video Recordings

The Camus system recorded individual videos from each home-cage with a frame resolution of 640×360 pixels, and video frame rate oof 20 frames per second. Our computer was built using a Thermaltake Toughpower GF3 1650W power supply, Asus PRIME A620-PLUS WIFI ATX AM5 motherboard, AMD Ryzen 9 7950X CPU, 32GB of Corsair Vengeance DDR5 RAM, and solid-state storage using a WD_BLACK SN850X NVMe SSD. To allow for 20-cameras to record videos simultaneously, we used five 4 Port USB 3.0 PCIe Card w/ 4 Dedicated 5Gbps Channels (USB 3.2 Gen 1) cards (StarTech PEXUSB3S44V) with 4 cameras plugged into each card. The computer was housed within a Raijintek Thetis Silver Classic case and mounted to the back of the cage rack.

The videos were saved using the XVID codec with an .avi container. Each frame for each camera was software timestamped according to the computer clock. The pixel to distance resolution of videos was calculated by video recording a sheet of paper with a 1cm-by-1cm grid placed at the bottom of the home cage. Following recording, pixels for each dimension were measured. We observed a mean value of 15.9 pixels/cm at the center for width and 16.5 pixels/cm at the center for height. The mean width values at horizontal edges were 14.3 pixels/cm and mean height values were 15.9 pixels/cm at vertical edges.

### Marker-less Keypoint Pose Estimation

For marker-less keypoint pose estimation, we used DeepLabCut (version 2.3.9).^29,49^ Specifically, we manually labeled 2,016 frames taken from 213 videos representing 44 different mice with either black or agouti coat colors. 95% of the dataset was used for training. The model was developed to extract 36 keypoints per frame from recorded videos. The list of extracted keypoints is provided in Supplementary Figure 1. We trained a ResNet-50 network initialized from ImageNet-pretrained weights, with location refinement (Huber loss, weight 0.05) and an output stride of 8. Optimization used stochastic gradient descent with a batch size of 8 and a piecewise-constant learning-rate schedule (0.005 for the first 10,000 iterations, 0.02 through 430,000, 0.002 through 730,000, and 0.001 thereafter). Images were augmented using the imgaug pipeline with rotation (±25°), scale jitter (0.5-1.25), cropping (crop ratio 0.4), CLAHE and histogram equalization, and emboss, each applied with probability 0.5. All other parameters were left at DeepLabCut defaults. Training ran for 950,000 iterations (∼87.5 h, TensorFlow), with network loss decreasing from 0.039 at 1,000 iterations to 0.0043 at 950,000 training iterations.

### Behavioral feature analysis

Behavioral features were extracted from DeepLabCut keypoints in MATLAB (R2023b), quantifying locomotor velocity, feeding, drinking, and nest-building for individually housed mice. Mouse position was computed as a body center per frame using the mean of major body keypoints detected with likelihood > 0.9 (Nosetip, EyeR, EyeL, EaroutsideR, EaroutsideL, Neck, CenterR, CenterC, CenterL, ThighR, ThighL, TailBase). Velocity was measured as the pixel distance this center point moved across time between frames, expressed as arbitrary units (au) as we did not convert pixel/time to cm/time, to acknowledge possible distortion at edges of images. Feeding and drinking were detected by the proximity of facial keypoints to apparatus landmarks, using thresholds chosen to maximize agreement with manual annotation as described below (Supplementary Figure 2).

Feeding was scored when the mouse head (mean of high likelihood nose, eye, ear, and neck keypoints) came within 41 pixels of the food-hopper axis, defined as the line segment between the North and South Food Hopper landmarks. Feeding behavior represents a complex set of embedded behaviors such as approach, sniff, bite, and chew with inherent variability across mice but are represented here as an interaction bout. Drinking was scored when the nose tip came within 21 pixels of the water-spout tip; but because the nose tip can be partially occluded while drinking, it was detected at a lower likelihood threshold (> 0.6) than other keypoints (> 0.9). Hopper and Spout landmarks were fixed at the median location of their confident detections (likelihood >0.9) to minimize keypoint jitter related errors. Feeding and drinking detection rules were validated against a three-rater consensus ground truth (agreement of ≥2 of 3 raters over ≥90% of each 1-s bin), pooled across videos manually annotated using BORIS.^50^ Thresholds were selected to maximize the Matthews correlation coefficient (Supplementary Figure 1).^51^ Extracted per-session features were exported in tabular (CSV) format for statistical analysis. All feature-extraction code and documentation are made available.

### Nest-building analysis

Nest-building was quantified with two complementary measures that track opposite ends of the same process: disappearance of the nestlet (Nestlet likelihood) and appearance of a cohesive nest (Nest likelihood). Nestlet likelihood was defined per frame as the DeepLabCut confidence likelihood detection of the nestlet keypoint. Nest likelihood was computed per frame as the mean confidence of eight keypoints placed around the nest perimeter in cardinal directions (North, NorthEast, East, SouthEast, South, SouthWest, West, NorthWest). Because these points are only detected confidently once a nest takes on its characteristic ring-like structure, their mean confidence rises as the nest is assembled and falls as it is disrupted, giving a graded index of nest formation.

We validated the utility of the calculated Nestlet likelihood and Nest likelihood against consensus based manual image annotations. At least two of three reviewers manually annotated each DeepLabCut extracted frame (2,127 frames from 212 recordings, representing 203 sessions from 41 different single mice and 9 sessions from 5 different pairs of mice) using a custom Matlab GUI. For each frame, reviewers scored nests using a modified 1-5 scale^24^ based on visual inspection of the top-down image. Reviewers were blinded to DeepLabCut based metrics when assigning nest scores. We computed Krippendorff’s alpha to assess preliminary interrater reliability, but all frames were also discussed to achieve consensus if there was any disagreement.

We computed the cumulative distributions of Nestlet likelihood and Nest likelihood values (each bounded between 0-1) for each ordinal level of consensus nest score (1-5). The cumulative distributions for each level were summarized as an area under the curve to compare values between levels.

To verify the circadian-like nature of nest building we also manually scored nests (Supplementary Figure 5) for 2 Males and 2 Females by assigning a number on the 1-5 scale per hour of the 72-hour recordings.

### Statistical Analysis for Camus Behavioral Features

All analyses were performed in R (version 4.5.3) using the lme4 and emmeans packages.^48,52^ For sex-specific circadian (window) analyses, each 72-hour session was partitioned into six Windows (dark and light windows of days 1-3) and their binned mean in each window modeled as **Value ∼ Sex × Window + (1 | Mouse/Session**), where the nested random effect captures both between-mouse variation and repeated within-mouse session variability. Omnibus effects were assessed with Type III tests of fixed effects. Post-hoc comparisons were computed with estimated marginal means (emmeans), using Sidak-adjusted pairwise contrasts on all 5 Window transitions (e.g., Day 1 Dark - Day 1 Light) with degrees of freedom by the Kenward-Roger method. A significant Sex × Window interaction was required before interpreting window-specific group differences. Post-hoc comparisons were computed with estimated marginal means (emmeans), using Holm-adjusted pairwise contrasts for effect of Sex on each Window (e.g., F-M in Day 1 Dark) with degrees of freedom by the Kenward-Roger method.

To examine sex-specific circadian transitions across windows, we split each window split into 6 hours windows, with a lead-in window ‘a’ and a terminal 6-hour window ‘b’ (e.g., Day 1 Dark ‘a’ and Dark ‘b’). The dozen six-hour windows were modeled as **Value ∼ Sex × Window + (1 | Mouse/Session**), where the nested random effect captures both between-mouse variation and repeated within-mouse session variability. Post-hoc comparisons were computed with estimated marginal means (emmeans) to contrast terminal 6-hour ‘b’ windows across successive windows within each sex (e.g., Day 1 Dark ‘b’ vs. Day 1 Light ‘b’) with Holm-adjustment and degrees of freedom by the Kenward-Roger method.

For aging analyses, each session was summarized as mean over the full 72-hour recording and modeled as **Value ∼ AgeBin + (1 | Mouse**), where the random intercept accounts for animals contributing sessions at multiple ages (ages binned into 2-6, 6-12, 12-18 months). WT and 5xFAD mice were modeled separately across whole sessions to avoid zero-one inflation related statistical artifacts in likelihood measures. Omnibus age effects were assessed with Type III tests of fixed effects, followed by Tukey-adjusted pairwise contrasts between age bins using estimated marginal means (emmeans), with degrees of freedom by the Kenward-Roger method.

## Acknowledgements

We wish to acknowledge Dr. Robert Vassar and Jelena Popovic for assistance with breeding the 5xFAD mice, and Tamra Whittenberg and the Center for Comparative Medicine at Northwestern University for their partnership in this project. We are also grateful for biostatistics advice from Dr. Kwang-Youn A Kim as part of our consultation with Biostatistics Collaboration Center at Northwestern University. We also want to thank Sebastián Malagón Pérez from the Kennedy Lab for his inputs on manually annotated behavioral analysis. We would also like to recognize the members of the Kimchi Lab for all their support and help in making this project successful.

## Author Contributions

TM, NA, GS & EK contributed to design of the work. TM, AL, SS, RP, VI, CYN, CNN & YG contributed to the acquisition of the data. TM & EK contributed to analysis and interpretation of data. TM, NA, & EK drafted the work. All authors reviewed the manuscript.

## Data Availability Statement

All code and design files presented in this manuscript are available from the Camus GitHub repository: https://github.com/KimchiLab/Camus/

## References

1. Tinbergen, N. On aims and methods of Ethology. Z. Für Tierpsychol. 20, 410–433 (1963).

2. Antoniadis, E. A., Ko, C. H., Ralph, M. R. & McDonald, R. J. Circadian rhythms, aging and memory. Behav. Brain Res. 114, 221–233 (2000).

3. Krakauer, J. W., Ghazanfar, A. A., Gomez-Marin, A., MacIver, M. A. & Poeppel, D. Neuroscience Needs Behavior: Correcting a Reductionist Bias. Neuron 93, 480–490 (2017).

4. Dennis, E. J. et al. Systems Neuroscience of Natural Behaviors in Rodents. J. Neurosci. 41, 911– 919 (2021).

5. Ding, S. S. et al. Fantastic beasts and how to study them: rethinking experimental animal behavior. J. Exp. Biol. 227, jeb247003 (2024).

6. K.S.V., A. B. G., Dissanayake, D., Gunatilake, M., Kuzhandai Velu, V. & Paranthaman, M. A short review on behavioural assessment methods in rodents. Bioinformation 19, 866–870 (2023).

7. Brown, L. A., Hasan, S., Foster, R. G. & Peirson, S. N. COMPASS: Continuous Open Mouse Phenotyping of Activity and Sleep Status. Wellcome Open Res. 1, 2 (2016).

8. Grieco, F. et al. Measuring Behavior in the Home Cage: Study Design, Applications, Challenges, and Perspectives. Front. Behav. Neurosci. 15, (2021).

9. Mösch, L. et al. Towards substitution of invasive telemetry: An integrated home cage concept for unobtrusive monitoring of objective physiological parameters in rodents. PloS One 18, e0286230 (2023).

10. Murphy, T. H. et al. High-throughput automated home-cage mesoscopic functional imaging of mouse cortex. Nat. Commun. 7, 11611 (2016).

11. Salem, G. et al. MouseVUER: video based open-source system for laboratory mouse home-cage monitoring. Sci. Rep. 14, 2662 (2024).

12. Tecott, L. H. & Nestler, E. J. Neurobehavioral assessment in the information age. Nat. Neurosci. 7, 462–466 (2004).

13. T, P., X, B., Mjh, K. & B, H. RFID-supported video tracking for automated analysis of social behaviour in groups of mice. *PubMed* https://pubmed.ncbi.nlm.nih.gov/31255597/ (2019).

14. Kahnau, P. et al. A systematic review of the development and application of home cage monitoring in laboratory mice and rats. BMC Biol. 21, 256 (2023).

15. Isik, S. & Unal, G. Frontiers | Open-source software for automated rodent behavioral analysis. 2023 https://doi.org/10.3389/fnins.2023.1149027 doi:10.3389/fnins.2023.1149027.

16. Luxem, K. et al. Open-source tools for behavioral video analysis: Setup, methods, and best practices. eLife 12, e79305 (2023).

17. De Visser, L., Van Den Bos, R., Kuurman, W. W., Kas, M. J. H. & Spruijt, B. M. Novel approach to the behavioural characterization of inbred mice: automated home cage observations. Genes Brain Behav. 5, 458–466 (2006).

18. Riedel, G. et al. Synthetic and plant-derived cannabinoid receptor antagonists show hypophagic properties in fasted and non-fasted mice. Br. J. Pharmacol. 156, 1154–1166 (2009).

19. Sharp, P. E., La Regina, M. C. & Suckow, M. A. The Laboratory Rat. (CRC Press, Boca Raton, 1998).

20. Ghosal, S. et al. Mouse handling limits the impact of stress on metabolic endpoints. Physiol. Behav. 150, 31–37 (2015).

21. Winter, Y. & Schaefers, A. T. U. A sorting system with automated gates permits individual operant experiments with mice from a social home cage. J. Neurosci. Methods 196, 276–280 (2011).

22. Singh, S., Bermudez-Contreras, E., Nazari, M., Sutherland, R. J. & Mohajerani, M. H. Low-cost solution for rodent home-cage behaviour monitoring. PLoS ONE 14, e0220751 (2019).

23. Salem, G. H. et al. SCORHE: a novel and practical approach to video monitoring of laboratory mice housed in vivarium cage racks. Behav. Res. Methods 47, 235–250 (2015).

24. Deacon, R. M. Assessing nest building in mice. Nat. Protoc. 1, 1117–1119 (2006).

25. Matikainen-Ankney, B. A. et al. An open-source device for measuring food intake and operant behavior in rodent home-cages. eLife 10, e66173 (2021).

26. Peirson, S. N., Brown, L. A., Pothecary, C. A., Benson, L. A. & Fisk, A. S. Light and the laboratory mouse. J. Neurosci. Methods 300, 26–36 (2018).

27. Mazzucato, L. Neural mechanisms underlying the temporal organization of naturalistic animal behavior. eLife 11, e76577 (2022).

28. Bedbrook, C. N. et al. Lifelong behavioral screen reveals an architecture of vertebrate aging.Science 391, eaea9795 (2026).

29. Mathis, A. et al. DeepLabCut: markerless pose estimation of user-defined body parts with deep learning. Nat. Neurosci. 21, 1281–1289 (2018).

30. Pereira, T. D. et al. SLEAP: A deep learning system for multi-animal pose tracking. Nat. Methods 19, 486–495 (2022).

31. Bains, R. S. et al. Assessing mouse behaviour throughout the light/dark cycle using automated in-cage analysis tools. J. Neurosci. Methods 300, 37–47 (2018).

32. Wang, C. et al. A comparative study of sleep and diurnal patterns in house mouse (Mus musculus) and Spiny mouse (Acomys cahirinus). Sci. Rep. 10, 10944 (2020).

33. Bailey, M. & Silver, R. Sex Differences in Circadian Timing Systems: Implications for Disease. Front. Neuroendocrinol. 35, 111–139 (2014).

34. Pastrick, A. et al. Biological Sex Influences Daily Locomotor Rhythms in Mice Held Under Different Housing Conditions. J. Biol. Rhythms 39, 351–364 (2024).

35. Freitag, N. et al. Sleep loss overrides sleep-preparatory behavior to promote sleep. Sleep https://doi.org/10.1093/sleep/zsag133 (2026) doi:10.1093/sleep/zsag133.

36. Jirkof, P. Burrowing and nest building behavior as indicators of well-being in mice. J. Neurosci. Methods 234, 139–146 (2014).

37. Gaskill, B. N. et al. Heat or Insulation: Behavioral Titration of Mouse Preference for Warmth or Access to a Nest. https://doi.org/10.1371/journal.pone.0032799 doi:10.1371/journal.pone.0032799.

38. Forner, S. et al. Systematic phenotyping and characterization of the 5xFAD mouse model of Alzheimer’s disease. Sci. Data 8, 270 (2021).

39. Oblak, A. L. et al. Comprehensive Evaluation of the 5XFAD Mouse Model for Preclinical Testing Applications: A MODEL-AD Study. Front. Aging Neurosci. 13, (2021).

40. Keszycki, R. et al. Characterization of apathy-like behaviors in the 5xFAD mouse model of Alzheimer’s disease. Neurobiol. Aging 126, 113–122 (2023).

41. O’Leary, T. P. & Brown, R. E. Age-related changes in species-typical behaviours in the 5xFAD mouse model of Alzheimer’s disease from 4 to 16 months of age. Behav. Brain Res. 465, 114970 (2024).

42. Reiber, M. et al. Development of behavioral patterns in young C57BL/6J mice: a home cage-based study. Sci. Rep. 12, 2550 (2022).

43. Deeming, D. C. Nest construction in mammals: a review of the patterns of construction and functional roles. Philos. Trans. R. Soc. B Biol. Sci. 378, 20220138 (2023).

44. Tagawa, N., Tsuneoka, Y. & Funato, H. Nest-building behavior in laboratory mice: Multifunctional roles and neural mechanisms. Neurosci. Biobehav. Rev. 178, 106377 (2025).

45. Mlost, J., Dawli, R., Liu, X., Costa, A. R. & Dorocic, I. P. Evaluation of unsupervised learning algorithms for the classification of behavior from pose estimation data. https://www.cell.com/patterns/abstract/S2666-3899(25)00085-6.

46. Oakley, H. et al. Intraneuronal β-Amyloid Aggregates, Neurodegeneration, and Neuron Loss in Transgenic Mice with Five Familial Alzheimer’s Disease Mutations: Potential Factors in Amyloid Plaque Formation. J. Neurosci. 26, 10129–10140 (2006).

47. Bachmanov, A. A., Reed, D. R., Beauchamp, G. K. & Tordoff, M. G. Food Intake, Water Intake, and Drinking Spout Side Preference of 28 Mouse Strains. Behav. Genet. 32, 435–443 (2002).

48. Bates, D., Mächler, M., Bolker, B. & Walker, S. Fitting Linear Mixed-Effects Models Using lme4. J. Stat. Softw. 67, 1–48 (2015).

49. Nath, T. et al. Using DeepLabCut for 3D markerless pose estimation across species and behaviors. Nat. Protoc. 14, 2152–2176 (2019).

50. Friard, O. & Gamba, M. BORIS: a free, versatile open-source event-logging software for video/audio coding and live observations. Methods Ecol. Evol. 7, 1325–1330 (2016).

51. Chicco, D. & Jurman, G. The Matthews correlation coefficient (MCC) should replace the ROC AUC as the standard metric for assessing binary classification. BioData Min. 16, 4 (2023).

52. Searle, S. R., Speed, F. M. & Milliken, G. A. Population Marginal Means in the Linear Model: An Alternative to Least Squares Means. Am. Stat. 34, 216–221 (1980).

