## Supplement for "Camus: a video-based framework for continuous behavioral monitoring in the mouse home-cage"

*Corresponding Author

#### Supplementary Figures

*Supplementary Figure 1: List and examples of keypoints labeled for identification using markerless pose estimation.*

| POINT | EXAMPLE | POINT | EXAMPLE | POINT | EXAMPLE |
|---|---|---|---|---|---|
| Nose | 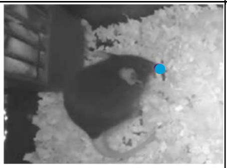 | Right Forepaw | 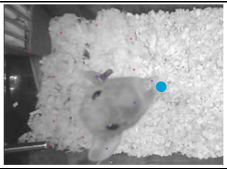 | Base Tail | 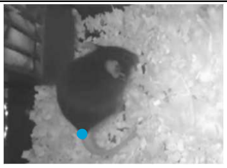 |
| Right Eye | 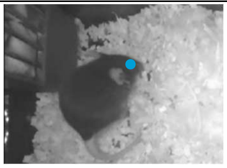 | Left Forepaw | 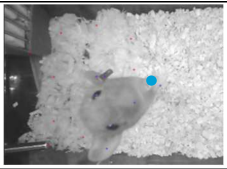 | Middle Tail | 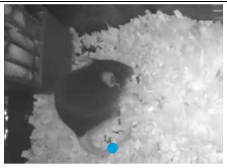 |
| Left Eye | 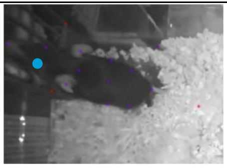 | Center right | 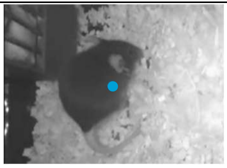 | Tip Tail | 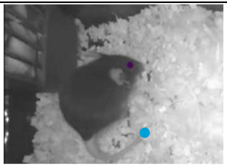 |
| Inner ear (right) | 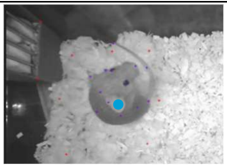 | Center | 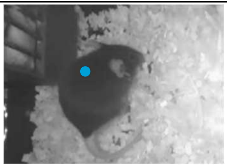 | Food Hopper North/South | 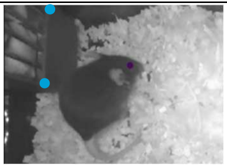 |
| Inner ear (left) | 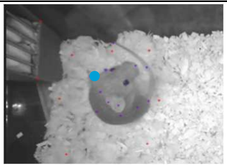 | Center left | 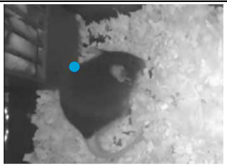 | Spout tip/base | 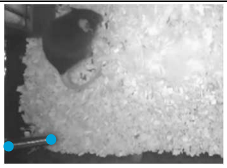 |
| Outer Ear (right) | 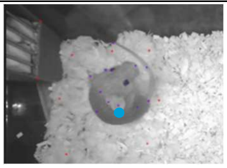 | Right Thigh | 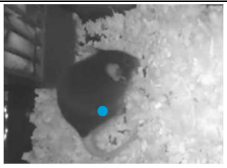 | Ventilation Hole | 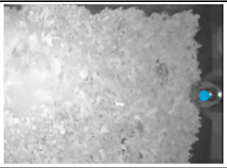 |
| Outer Ear (left) | 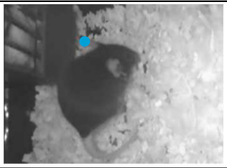 | Left Thigh | 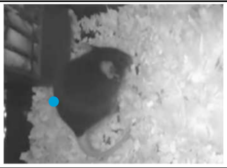 | Nestlet Center | 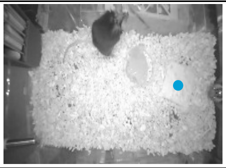 |
| Ear Tag (right or left depending on mouse) | 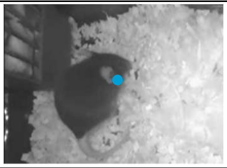 | Right Hind paw | 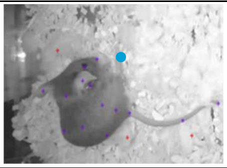 | Nest Cardinal Points (North, Northeast, East, Southeast, South, Southwest, West, Northwest) | 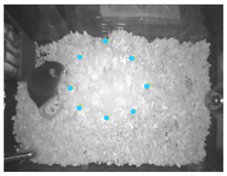 |
| Neck | 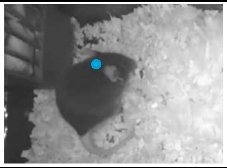 | Left Hind paw | 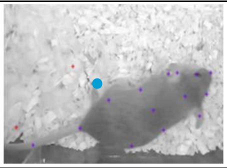 | | |

We used a combination of body-skeleton based keypoints as well as environmental keypoints. We illustration each keypoint with an example of the keypoint labeled on mice. Both agouti and black coated mice are shown in the examples. We also trained the model to label the center of the compressed Nestlet, and the cardinal directions of the formed Nest points that appear as the nestlet is shredded into a cohesive nest. We use cardinal direction points to determine the edges of the circular crater like nests and track the boundary of the nest as it was built.

*Supplementary Figure 2*

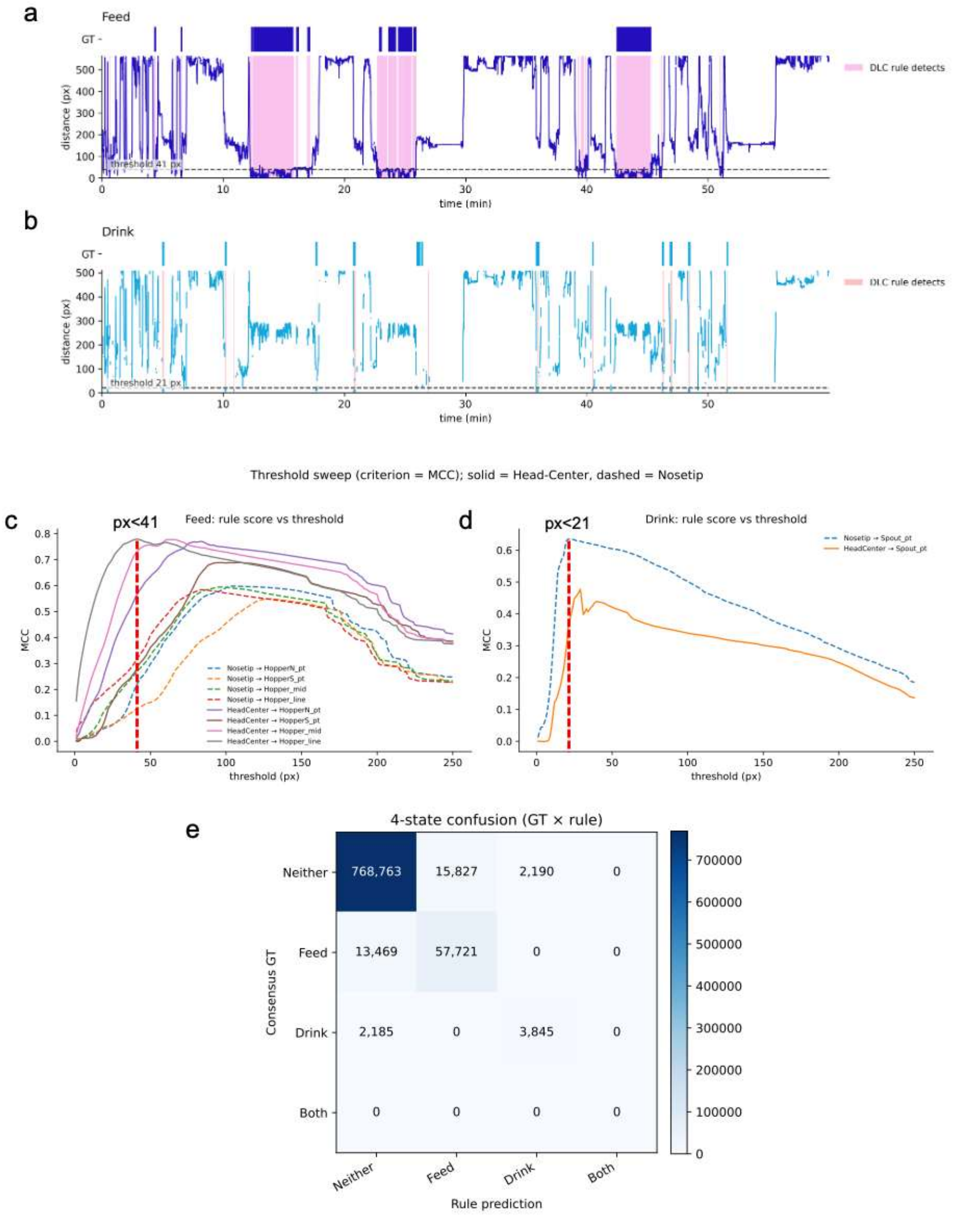


**a- b.** Feeding and Drinking behavior were detected from DeepLabCut keypoints using a distance threshold between mouse keypoints and the relevant apparatus feature, with the threshold optimized against manually annotated ground truth by Matthew's correlation coefficient (MCC). Representative traces of the mouse-to-apparatus distance over time are displayed for **a.** Feeding (dark blue; Mouse Head-to-Food Hopper distance) and **b.** Drinking (light blue; Nose tip-to-Waterspout distance). Dashed line indicates the selected distance threshold (Feed 41 pixels; Drink 21 pixels); shaded intervals mark periods the DeepLabCut rule classifies as detections (Feed, pink;

Drink, orange). Consensus ground-truth annotations are shown as the top raster in each panel. Detections correspond to intervals where the distance falls below threshold.

**c.** Threshold sweep of detection performance (MCC vs. distance threshold) across candidate keypoint pairings, comparing Head-Center (solid) and Nose tip (dashed) reference points against alternative apparatus rules. Red dashed line marks the selected threshold. For Feed, the Head-Center-to-Food Hopper pairing peaked near 41 pixels (peak MCC ≈ 0.8).

**d.** For Drink, the Nose tip-to-Spout pairing outperformed Head-Center and peaked near 21 pixels (peak MCC ≈ 0.63). The lower peak MCC for Drink reflects its shorter, sparser events and greater keypoint drop out at the spout.

**e.** Four-state confusion matrix (Neither / Feed / Drink / Both) between consensus Ground-truth (rows) and rule prediction (columns), pooled across sessions. The rule recovers most of the Feed and Drink events with errors concentrated in the Neither↔Feed or Drink boundary; Feed and Drink are never mutually confused, and the empty "Both" row/column confirms the two behaviors are non-overlapping.

*Supplementary Figure 3: Validation of automated nest measures against manual rater nest scoring using the 1-5 scale*

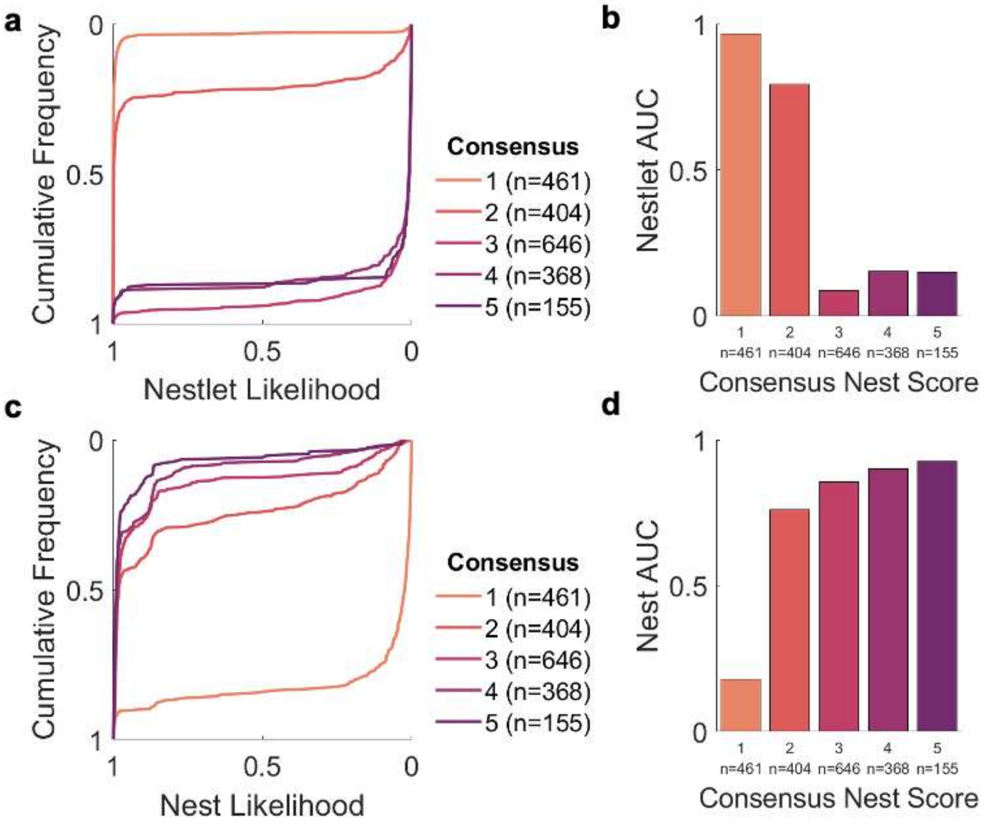


**a.** Nestlet likelihood: We validated the utility of the DeepLabCut derived Nestlet likelihood against consensus based manual image annotations. We computed the cumulative distributions (y-axis) of Nestlet likelihoods (x-axis) for each ordinal level of consensus nest score (1-5, colored curves). N values indicate the number of frames rated for each ordinal nest score level.

**b.** Cumulative distributions for each nest score level were summarized as an area under the curve (AUC) to compare values between ordinal nest score levels. DeepLabCut derived Nestlet likelihood disappearance tracked the shredding of the nestlet, consistent a transition from level 2 to 3 in the original scale.[24]

**c.** Nest likelihood: Cumulative distributions of DeepLabCut derived Nest likelihood against consensus based manual image annotations. Conventions as in **a.**

**d.** Summary AUC values for Nest likelihood by consensus nest scores. The distributions of Nest likelihood displayed a monotonic relationship with consensus nest scores.

*Supplementary Figure 4: Hourly nest scores labelled manually over 72 hours*

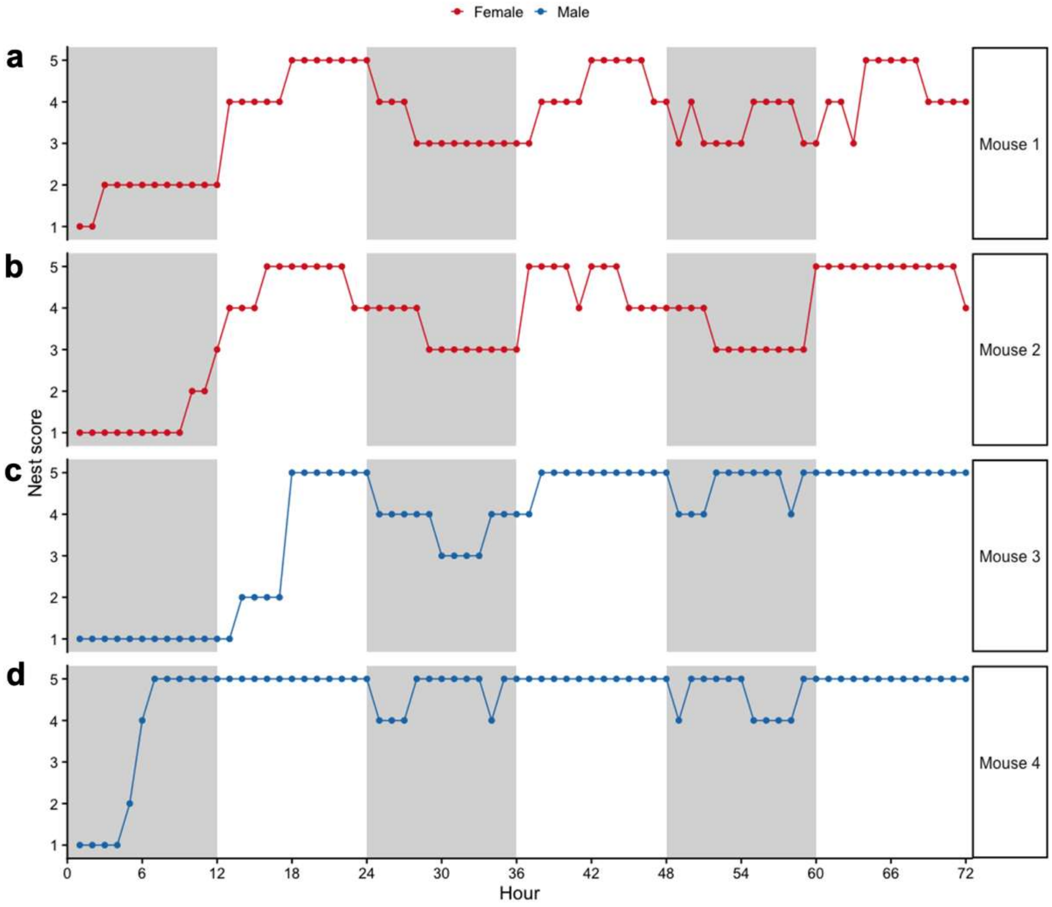


**a-d.** Nest scores were rated manually each hour on a 1-5 scale[24] over 72-hours across 4 mice (**a-b**. Females; **c-d**. Males). The graphs show variability in nest scores over time showing the dynamic nature of nest building, including frequent disruptions, especially during dark periods in female mice.

*Supplementary Figure 5: Sex-difference in nesting behavior during dark-light circadian window changes*

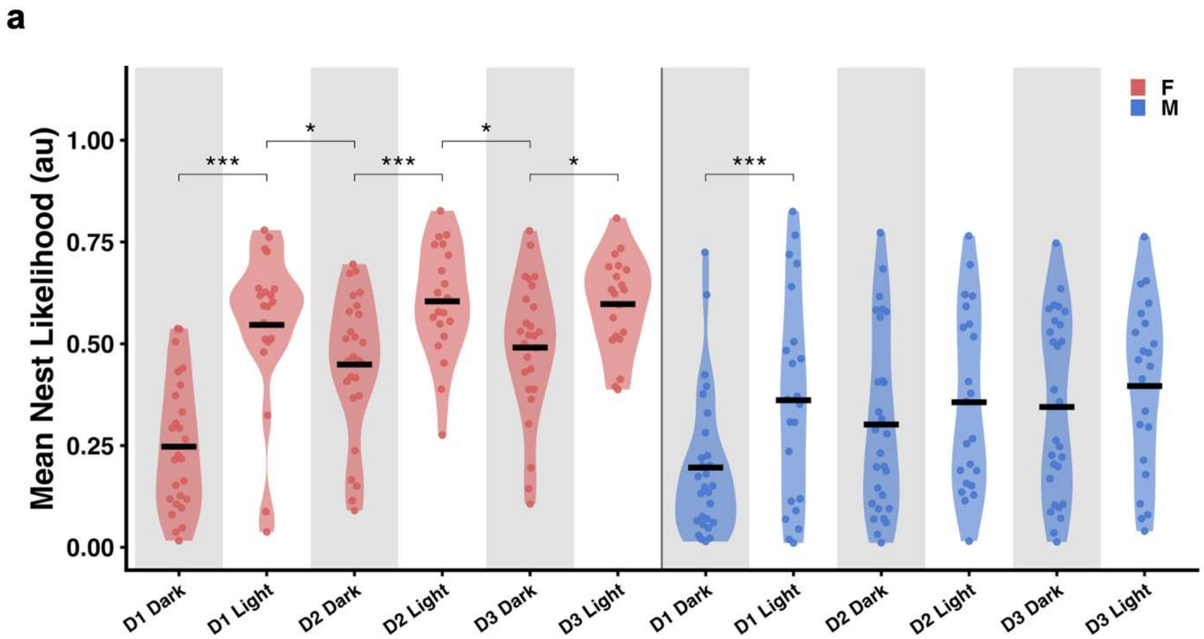


**a.** Mean Nest likelihood per 72 hour session, computed over the final 6 hour 'b' windows of each dark and light window (Day 1-Day 3), for female (red) and male (blue) WT mice (n = 27 and 29 sessions). The LMEM with all 12 Windows (Each window divided into 'a' and 'b'; e.g., Day 1 Dark 'a' and Dark 'b') showed a significant effect of Sex × Window interaction (LMEM, p=0.0002). Female mice consistently alternated Nest likelihoods across every window transition, with Nest likelihood rising across each dark-to-light 'b' transition (emmeans contrasts holm-adjusted, Day 1, $p < 0.0001$; Day 2, p =0.0009; Day 3, p = 0.0386) and falling across each light-to-dark 'b' transition (emmeans contrasts holm-adjusted, Day 2, p = 0.0386; Day 3, p = 0.0344). Male mice showed only an initial rise from dark-to-light 'b' transition (emmeans contrasts holm-adjusted, Day 1, $p < 0.0001$). Significance code for post-hoc tests: *** <0.001, ** <0.01, * <0.05 are represented with overlying comparison brackets.

#### Supplementary Tables

*Supplementary Table 1: DEEPLABCUT Model Training and Testing Errors over Iterations*

| | Training iterations: | Training dataset (%) | Shuffle number | Train error(pixels) | Test error (pixels) | p-cutoff used | Train error with p-cutoff | Test error with p-cutoff |
|---|---|---|---|---|---|---|---|---|
| 0 | 50000 | 95 | 7 | 17.27 | 18.73 | 0.6 | 7.69 | 8.12 |
| 1 | 100000 | 95 | 7 | 10.72 | 13.95 | 0.6 | 6.78 | 8.87 |
| 2 | 150000 | 95 | 7 | 8.47 | 13.12 | 0.6 | 6.35 | 8.99 |
| 3 | 200000 | 95 | 7 | 6.65 | 12.01 | 0.6 | 5.71 | 8.86 |
| 4 | 250000 | 95 | 7 | 5.43 | 11.77 | 0.6 | 4.88 | 9.35 |
| 5 | 300000 | 95 | 7 | 4.69 | 11.73 | 0.6 | 4.33 | 8.98 |
| 6 | 350000 | 95 | 7 | 4.32 | 11.84 | 0.6 | 4.04 | 9.39 |
| 7 | 400000 | 95 | 7 | 3.65 | 11.52 | 0.6 | 3.42 | 9.03 |
| 8 | 450000 | 95 | 7 | 3.41 | 11.42 | 0.6 | 3.14 | 9.22 |
| 9 | 500000 | 95 | 7 | 3.35 | 11.56 | 0.6 | 3.11 | 9.23 |
| 10 | 550000 | 95 | 7 | 3.39 | 11.68 | 0.6 | 3.14 | 9.38 |
| 11 | 600000 | 95 | 7 | 3.26 | 11.7 | 0.6 | 3.02 | 9.37 |
| 12 | 650000 | 95 | 7 | 3.32 | 11.53 | 0.6 | 3.05 | 9.26 |
| 13 | 700000 | 95 | 7 | 3.47 | 11.78 | 0.6 | 3.24 | 9.46 |
| 14 | 750000 | 95 | 7 | 3.25 | 11.65 | 0.6 | 3.01 | 9.27 |
| 15 | 800000 | 95 | 7 | 3.15 | 11.67 | 0.6 | 2.91 | 9.24 |
| 16 | 850000 | 95 | 7 | 3.21 | 11.71 | 0.6 | 2.97 | 9.31 |
| 17 | 900000 | 95 | 7 | 3.29 | 11.66 | 0.6 | 3.06 | 9.32 |
| 18 | 950000 | 95 | 7 | 3.18 | 11.64 | 0.6 | 2.95 | 9.33 |

### Supplementary Videos

*Supplementary Video 1 Recording of an individually housed mouse in Camus*

*Supplementary Video 2: Recording demonstrating 20 simultaneous Camus sessions*

*Supplementary Video 3 Video with DeepLabCut generated keypoint annotations*
